# Genetic contributions to trail making test performance in UK Biobank

**DOI:** 10.1101/103119

**Authors:** Saskia P Hagenaars, Simon R Cox, W David Hill, Gail Davies, David CM Liewald, CHARGE consortium Cognitive Working Group, Sarah E Harris, Andrew M McIntosh, Catharine R Gale, Ian J Deary

**Author notes:** These authors contributed equally to the work (Catharine R Gale, Ian J Deary). Corresponding author, Ian J. Deary, Centre for Cognitive Ageing and Cognitive Epidemiology, Department of Psychology, University of Edinburgh 7, George Square, Edinburgh, EH8 9JZ, Scotland, UK.

## Abstract

The Trail Making Test is a widely used test of executive function and has been thought to be strongly associated with general cognitive function. We examined the genetic architecture of the trail making test and its shared genetic aetiology with other tests of cognitive function in 23 821 participants from UK Biobank. The SNP-based heritability estimates for trail-making measures were 7.9 % (part A), 22.4 % (part B), and 17.6 % (part B – part A). Significant genetic correlations were identified between trail-making measures and verbal-numerical reasoning (r_g_ > 0.6), general cognitive function (r_g_ > 0.6), processing speed (r_g_ > 0.7), and memory (r_g_ > 0.3). Polygenic profile analysis indicated considerable shared genetic aetiology between trail making, general cognitive function, processing speed, and memory (standardized β between 0.03 and 0.08). These results suggest that trail making is both phenotypically and genetically strongly associated with general cognitive function and processing speed.

## Introduction

The Trail Making Test (TMT) is widely used in both research and clinical settings as a test of some aspects of executive function ^1-3^. The TMT is usually given as two parts, from which three measures are derived. In Part A (TMT-A), participants are required to connect an array of numbers in ascending order, by drawing a continuous line (trail) between them as quickly and accurately as possible. Part B (TMT-B) requires participants to connect an array of both numbers and letters in *alternating* ascending order (1, A, 2, B, 3, C, etc.) with the same emphasis on speed and accuracy (Supplementary Figure 1). Subtracting TMT-A completion time from that of TMT-B (TMT B minus A) is thought to allow the relative contributions of visual search and psychomotor speed to be parsed from the more complex executive functions (such as cognitive flexibility) required to alternate between numbers and letters ^4-8^.

**Figure 1.**
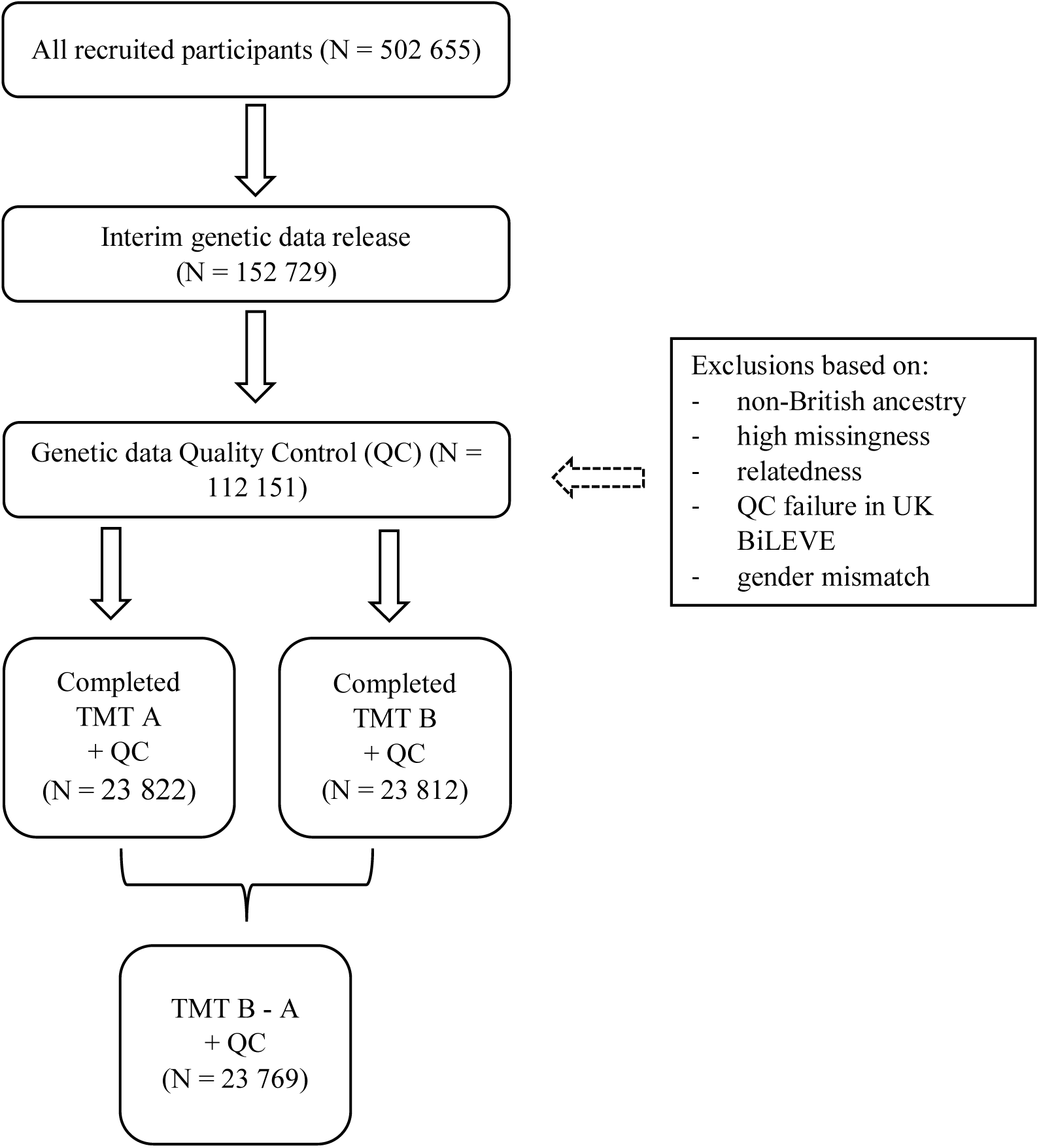
Flow diagram of participant selection

TMT performance has been ascribed to a number of cognitive processes, “including attention, visual search and scanning, sequencing and shifting, psychomotor speed, abstraction, flexibility, ability to execute and modify a plan of action, and ability to maintain two trains of thought simultaneously” (Salthouse et al., 2011, p222^9^). It is considered a useful tool in research and clinical practice due to the sensitivity of the task (particularly TMT B and B-A) to frontal lobe damage (in some, but not other studies; see MacPherson, Della Sala ^10^) and dementia ^11-13^. There are declines in both TMT A and B performance in ageing ^10, 14-20^. There is also evidence for performance deficits on TMT B in mood disorders ^21^, and in patients with schizophrenia and their relatives ^18, 22-27^

Family-based and twin-based studies have provided evidence for a genetic contribution to individual differences in trail making, estimating the heritability for trail making part A between 0.23 and 0.38, and between 0.39 and 0.65 for trail making part B ^28-30^. A recent genome-wide association study (GWAS) of trail making part A and part B in a sample of around 6000 individuals did not find any genome-wide significant hits ^31^; however, GWAS of other cognitive phenotypes have demonstrated that much larger sample sizes are required to reliably identify significant genetic loci ^32, 33^. Trail making is thought to have genetic influences that are shared with other cognitive abilities, with a twin-based genetic correlation of 0.48 reported between trail making, measured as the ratio between trail making part A and trail making part B, and general cognitive function, and 0.52 with working memory ^34^.

In addition to a relatively poor understanding of the molecular genetic underpinnings of TMT, its cognitive and psychometric architecture merits further research. Specifically, it is unclear whether the cognitive abilities required for TMT-B performance are distinct from other cognitive domains, because TMT A and B scores correlate with measures of general cognitive function and processing speed (correlation coefficient estimates range from ~0.3 to ~0.7; ^9, 20^) ^30, 35-38^. Evidence from cognitive-ageing studies further suggests a strong overlap between TMT performance and other cognitive domains, because age-related decline in processing speed and working memory account for much of the age effects on TMT-B ^9, 20, 39, 40^. However, reliably identifying the cognitive processes that underpin cognitive test performance using phenotypic correlational analyses alone is sub-optimale.g. ^41, 42, 43^. Rather, the interpretation of relationships between TMT performance and other cognitive abilities such as general cognitive function, may be enhanced by considering and comparing their respective shared and unshared causes, including their genetic architecture.

The aim of the present study is to: 1) add to the understanding of the genetic architecture of trail making; and 2) to explore the overlap between genetic architecture of TMT performance and other cognitive abilities, including general cognitive function. Although several studies have examined the genetic architecture of the trail making test and its overlap with other cognitive abilities, they had relatively low power. The largest TMT GWAS (N ~ 6000) to date used a variety of assessments across multiple cohorts across multiple countries, potentially leading to heterogeneity in both sample composition and in the cognitive measures ^31^. The current study design, using >23 000 UK Biobank participants, mitigates such confounds: the almost four-fold increase in sample size yields greater statistical power, and the single sample reduces heterogeneity in genetic testing and in phenotype, because all participants had white British ancestry and took the same TMT test with the same instructions. In addition, we add to this significantly larger GWAS than has previously been conducted with: (1) estimates of the SNP-based heritability of TMT performance, and (2) an examination of genetic overlap among the different TMT test measures and with other cognitive functions, using polygenic profile analysis and genetic correlations in independent samples.

## Materials and methods

### Participants

UK Biobank is a large resource which aims to identify determinants of human diseases in middle aged and older individuals (http://www.ukbiobank.ac.uk)^44^. A total of 502 655 community-dwelling individuals aged between 37 and 73 years were recruited in the United Kingdom between 2006 and 2010. Baseline assessment included cognitive testing, personality self-report, and physical and mental health measures. For the present study, genome-wide genotyping data were available for 112 151 participants (58 914 females, 53 237 males) after quality control (see below). They had a mean (SD) age of 56.9 (7.9) years (range 40 to 73 years). UK Biobank received ethical approval from the Research Ethics Committee (REC reference for UK Biobank is 11/NW/0382). This study was completed under UK Biobank application 10279. Figure 1 shows the study flow for the present study.

## Measures

### Trail making test

The trail making test parts A (TMT A) and B (TMT B) were introduced at a follow up testing wave in UK Biobank, between 2014 and 2015. For TMT A, participants were instructed to connect numbers (1 – 25) consecutively (which were quasi-randomly distributed on the touchscreen) as quickly as possible in ascending order by selecting the next number. TMT B is similar, but letters (A – L) and numbers (1 – 13) had to be selected in alternating ascending order, e.g. 1 A 2 B 3 C etc. The intervals between touching two points was timed in seconds using a Javascript timer. The total time (in seconds) to complete the trail making test (part A or B) was derived by summing the interval values between two points. Nine out of 23 821 individuals who scored >250 seconds for TMT B were excluded. Owing to positively skewed distributions, both TMT A and TMT B scores were log-transformed prior to further analyses. The difference between the raw scores for TMT A and TMT B was computed as TMT B minus TMT A (TMT B-A). 52 out of 23 821 individuals with scores <-50 or >150 were removed from TMT B-A. After exclusions, 23 822 individuals with genetic data completed TMT A, 23 812 individuals with genetic data completed TMT B, and 23 769 individuals had complete information for TMT B-A. The three trail making measures have been scored such that a higher score indicates better performance. This study also used the verbal-numerical reasoning test from UK Biobank (VNR, N = 36 035), which consisted of a 13-item questionnaire assessing verbal and arithmetical deduction as described by Hagenaars et al.^45^.

### Genotyping and quality control

The interim release of UK Biobank included genotype data for 152 729 individuals, of whom 49 979 were genotyped using the UK BiLEVE array and 102 750 using the UK Biobank axiom array. These arrays have over 95% content in common. Details of the array design, genotyping procedures and quality control details have been published elsewhere ^45, 46^. UK Biobank released an imputed dataset as part of the interim data release. More details can be found at the following URL: http://biobank.ctsu.ox.ac.uk/crystal/refer.cgi?id=157020. A minor allele frequency cut-off of 0.1 % was used for all autosomal variants, as well as an imputation quality score above 0.1 (N ~ 17.3M SNPs).

### Genome-wide association analysis (GWAS)

Genotype-phenotype association analyses were conducted using SNPTEST v2.5.1 ^47^. The ‘frequentist 1’ option was used to specify an additive model, and genotype dosages were analysed to account for genotype uncertainty. All phenotypes were adjusted for the following covariates prior to analysis; age, gender, assessment centre, genotyping array and batch and the first 10 genetic principal components for population stratification. The GWAS of VNR has been performed previously ^32^.

### SNP-based heritability and genetic correlations

Univariate GCTA-GREML ^48^ analysis was performed to estimate the proportion of variance explained by all genotyped common SNPs for TMT A, TMT B and TMT B-A. To include only unrelated individuals, a relatedness cut-off of 0.025 was used in the generation of the genetic relationship matrix. Bivariate GCTA-GREML ^48^ and LD score regression ^49^ were used to derive genetic correlations between TMT measures and VNR in UK Biobank ^32^. LD score regression was also used to estimate genetic correlations between trail making measures in UK Biobank and trail making part A, trail making part B, general cognitive function, processing speed, and memory from the CHARGE consortium meta-analyses of these cognitive phenotypes ^31, 33, 50^.

### Gene-based association analysis

MAGMA ^51^ was used to derive gene-based associations using the summary results of the three GWAS for trail making. 18 062 genes were analysed after the SNPs were assigned a gene based on their position using the NCBI 37.3 build, without additional boundaries placed around the genes. To account for linkage disequilibrium, the European 1000 Genomes data panel (phase 1, release 3) was used as a reference. A Bonferroni correction was used to control for 18 062 tests (α= 0.05/18 062; P < 2.768 x 10^-6^). The gene-based associations for trail making were compared with gene-based associations for VNR, and with the gene-based associations for trail making, general cognitive function, processing speed, and memory from the CHARGE consortium, based on the GWAS summary results ^31, 33, 50^. The CHARGE summary results were converted from HapMap2 to 1000G format to ensure the maximum overlap between the two samples. This was achieved using the LiftOver program, which converts coordinate ranges between genome assemblies.

### Partitioned heritability

Partitioned heritability analyses were performed on the trail making SNP-based association results to determine if SNPs group together according to a specific biological function or role and thereby making an enriched contribution to the total proportion of heritability of the trail making phenotypes. These analyses were performed using the data processing pipeline as suggested by Finucane et al. ^52^.

### Polygenic profile analyses

Polygenic profile analyses were performed to predict trail making test performance into UK Biobank and to predict cognitive function scores in two independent cohorts, Generation Scotland’s Scottish Family Health Study (GS) and the Lothian Birth Cohort of 1936 (LBC1936). Prediction from the CHARGE consortium meta-analysis of trail making, general cognitive function, processing speed, and memory into UK Biobank was performed to test the extent to which individual differences in trail making tests in UK Biobank could be predicted by the polygenic architecture of these four traits.

#### Polygenic prediction into UK Biobank

The UK Biobank genotyping data were recoded from numeric (1,2) allele coding to standard ACGT coding using a bespoke programme developed by one of the present authors (DCL) ^45^. Polygenic profile scores were created for trail making part A, trail making part B, general cognitive function, processing speed, and memory based on the results from the CHARGE consortium meta-analysis in all genotyped participants using PRSice ^53^. SNPs with a minor allele frequency < 0.01 were removed prior to creating the scores. Clumping was used to obtain SNPs in linkage disequilibrium with an r2 < 0.25 within a 250kb window. Five polygenic profiles were created for each of the three phenotypes according to the significance of the association with the trail making phenotype, at p-value thresholds of 0.01, 0.05, 0.1, 0.5 and 1 (all LD pruned SNPs). Regression models were used to examine the association between the polygenic profiles and TMT A, TMT B and TMT B-A phenotype scores in UK Biobank, adjusting for age at measurement, sex, genotyping batch and array, assessment centre, and the first ten genetic principal components for population stratification. All associations were corrected for multiple testing using the false discovery rate (FDR) method ^54^.

#### Polygenic prediction into GS and LBC1936

Polygenic profile scores were created in PRSice ^53^, using the UK Biobank trail making SNP-based association results, for genotyped participants of GS (n = 19 994) and LBC1936 (n = 1005). Individuals were removed from GS if they had contributed to both UK Biobank and GS (n = 174). Polygenic profile scores were created based on the significance of the association in UK Biobank with the trail making phenotype, at p-value thresholds of 0.01, 0.05, 0.1, 0.5 and 1 (all SNPs). Linear regression models were created to test the association between the polygenic profiles for trail making and the target phenotypes in: GS (Wechsler digit-symbol substitution ^55^, phonemic verbal fluency ^1^, Wechsler logical memory ^56^, the Mill Hill vocabulary test ^57^, a fluid cognitive function component, and a general cognitive function component); and LBC1936 (trail making part B, a fluid cognitive function component, vocabulary, memory, processing speed, change in fluid cognitive function between age 11 and age 70, Moray House Test score at age 11, and Moray House Test score at age 70). Further details about the cognitive tests can be found in the Supplementary Materials. All models were adjusted for age, sex and the first five (GS) or four (LBC1936) principal components for population stratification. The GS models were also adjusted for family structure by fitting a univariate linear mixed model which estimates the genetic and environmental variance, using the ASReml program ^58^. All associations were corrected for multiple testing using the FDR method ^54^.

## Meta-analysis

Inverse variance-weighted meta-analysis of UK Biobank trail making and CHARGE trail making GWAS was performed using the METAL package (http://www.sph.umich.edu/csg/abecasis/Metal). The meta-analysis was restricted to SNPs that were available in both samples (N SNPs TMT A = 2 332 746; N SNPs TMT B = 2 466 810), and the samples did not include any overlapping individuals. The total sample in the meta-analysis consisted of 29 251 individuals for TMT A and 30 022 for TMT B.

## Results

### Phenotypic correlations

23 822 individuals with genetic data completed TMT A, 23 812 individuals with genetic data completed TMT B, and 23 769 individuals with genetic data had complete information for TMT B-A. Table 1 shows the phenotypic correlations between the TMT phenotypes, as well as with verbal-numerical reasoning in UK Biobank. Correlations indicated that individuals who took more time to complete the trail making tests had lower performance on the verbal-numerical reasoning test. The strongest correlation between VNR and TMT was found for TMT B (0.36). Supplementary Figure 2 shows the age and sex distribution of the different trail making measures.

**Table 1.** Phenotypic correlations for the UK Biobank cognitive tests in all genotyped participants. Standard errors for the correlations are shown in parentheses. Pearson correlations were used for continuous-continuous correlations for the phenotypic correlations.

|  | TMT A | TMT B | TMT B-A | VNR |
| --- | --- | --- | --- | --- |
| TMT A | - |  |  |  |
| TMT B | 0.62 (0.004) | - |  |  |
| TMT B-A | 0.04 (0.006) | 0.80 (0.002) | - |  |
| VNR | 0.20 (0.010) | 0.36 (0.009) | 0.32 (0.010) | - |

TMT A, trail making part A; TMT B, trail making part B; TMT B-A, trail making part B – part A; VNR, verbal-numerical reasoning.

**Figure 2.**
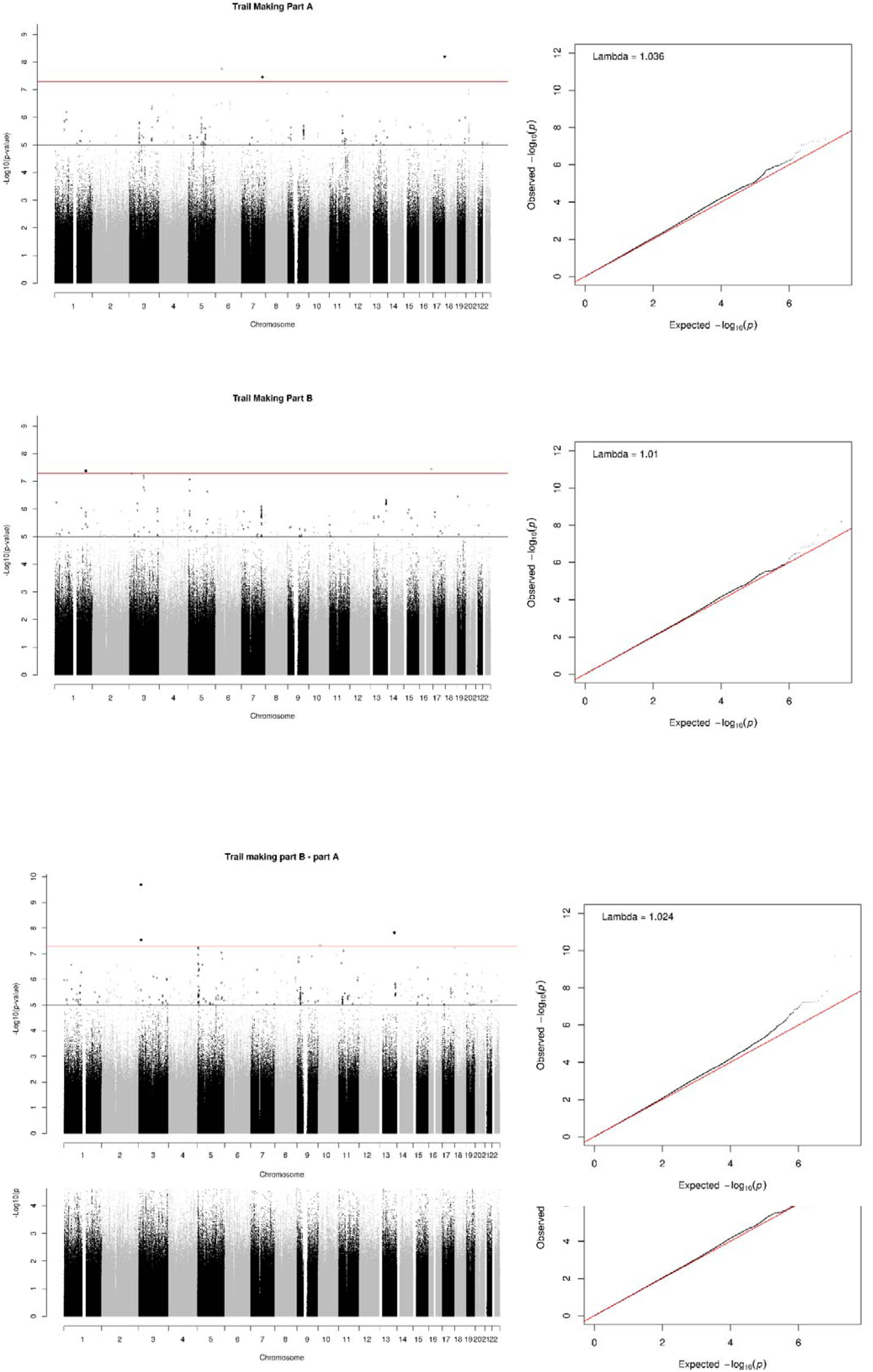
Manhattan and Q-Q plot of P-values of the SNP-based association analysis of Trail Making Part A, Trail Making Part B and Trail Making Part B – Part A. The red line in the Manhattan plots indicates the threshold for genome-wide significance (P < 5 × 10^-8^); the grey line indicates the threshold for suggestive significance (P < 1 × 10^-5^).

### Genome-wide association study

The results of the GWAS analyses are presented in Figure 2; for each trail making phenotype a Manhattan and a QQ plot is shown.

#### Trail making test part A

For TMT A three SNPs reached genome-wide significance. These were located in three loci, on chromosome 6, 7 and 17. The strongest signal was on chromosome 17 (Affymetrix id 17:77673843_T_C; p = 6.31 x 10^-9^, MAF = 0.0016); this SNP is not in a gene and does not have an rs id. The top SNP on chromosome 6 (rs185657300, p = 1.77 x 10^-8^, MAF = 0.0013) is also not located within a gene. The top SNP on chromosome 7 (rs554959089, p = 3.49 x 10^-8^, MAF = 0.0020) is located in a region that includes the Homeodomain Interacting Protein Kinase 2 (*HIPK2*) gene, a tumour suppressor. Activation of *HIPK2* leads to an apoptotic response after DNA damage in cells ^59^. None of the three top SNPs were observed in large association peaks and all had low minor allele frequencies; therefore, these associations should be treated with caution until replication is observed in an independent sample.

Gene based analyses identified three genes that were significantly associated with TMT A (Table 2 and Supplementary Table 1a); *CRNKL1*, a protein necessary for pre-mRNA splicing on chromosome 20 ^60^; caspase 5 (*CASP5)*, which plays a role in the execution phase of cell apoptosis on chromosome 11 ^61^; and *NAA20*, a component of the N-acetyltransferase complex B on chromosome 20 ^62^. All three of these genes were also nominally significant (p < 0.05) in the GWAS of TMT B, and *CASP5* was also nominally significant in the analyses for CHARGE general cognitive function and processing speed (Table 2).

**Table 2.**
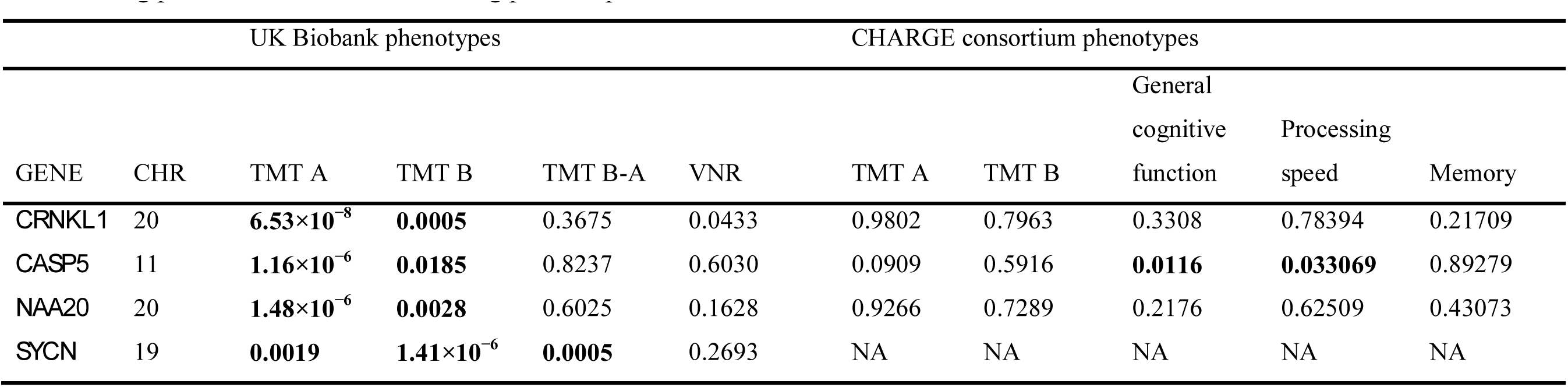
Shows the genome wide significant genes (P < 2.768 × 10^-6^) from the UK Biobank trail making phenotypes and the significance values for the same genes using the other two trail making phenotypes and Verbal Numerical Reasoning (VNR) measured in UK Biobank, and cognitive phenotypes from the CHARGE consortium. Unmodified P-values are shown for all phenotypes. TMT A, trail making part B – part A.

The proportion of variance in TMT A explained by all common genetic variants using GCTA-GREML was 0.079 (SE 0.024).

#### Trail making test part B

Two SNPs exhibited genome-wide significance for TMT B, on chromosome 1 (rs34804445, p = 4.18 x 10^-8^, MAF = 0.0253) and chromosome 16 (rs541566700, p = 3.59 x 10^-8^, MAF = 0.0034). Neither SNP was located within a gene. Gene based analyses identified one gene (*SYCN* on chromosome 19) associated with TMT B (Table 2 and Supplementary Table 1b); this gene is expressed in the human pancreas ^63^. This gene was also nominally significant (p < 0.05) in the gene-based analysis for the other trail making variables measured in UK Biobank. Results for this gene were not available for the CHARGE consortium phenotypes (Table 2).

The proportion of variance in TMT B explained by all common genetic variants using GCTA-GREML was 0.224 (SE 0.026).

#### Trail making test part B – part A

Five SNPs reached genome-wide significance for TMT B-A spanning three regions on chromosome 3, 10 and 13. The two genome-wide significant SNPs on chromosome 3 were not located within genes. The top SNP on chromosome 10 (rs549730795, p = 4.81 x 10^-8^, MAF = 0.0028) is located in a region that includes *CAMK1D*, calcium/calmodulin-dependent protein kinase 1 family ^64^. One genome-wide significant SNP, rs529887265 (p = 1.50 x 10^-8^, MAF = 0.0013), was observed on chromosome 13; this region includes the *DOCK9* gene, which is part of the evolutionary conserved exchange factors for the Rho GTPases ^65^. Gene-based analyses did not identify any significant associations for TMT B-A (Supplementary Table 1c).

The proportion of variance in TMT B-A explained by all common genetic variants using GCTA-GREML was 0.176 (SE 0.025).

### Partitioned heritability

Significant enrichment was found for TMT B in evolutionary conserved regions with a 500bp boundary, where 33 % of the SNPs accounted for 95 % of the heritability (enrichment metric = 2.85, SE = 0.44, p = 2.42 × 10^−5^). TMT A and TMT B-A were unsuitable for this analysis, due to low heritability Z-scores, which were 2.6 and 5.4 respectively ^52^.

### Genetic correlations

The results of the bivariate GCTA-GREML and LD score regression analyses within UK Biobank are shown in Table 3. The LD score regression analyses with the cognitive phenotypes from the CHARGE consortium are shown in Table 4. Strong positive genetic correlations, using GCTA-GREML, were observed between all three measures derived from the trail making test in UK Biobank (r_g_ between 0.64 and 0.96). Large genetic correlations were found between the three trail making tests and verbal-numerical reasoning (r_g_ between 0.59 and 0.64). Similar results were found when calculating the genetic correlations using LD score regression (Table 3). Positive genetic correlations were observed between trail making in UK Biobank and the following GWAS meta-analyses from the CHARGE consortium: general cognitive function (r_g_ between 0.61 and 0.70), processing speed (r_g_ between 0.69 and 0.76), and memory (r_g_ between 0.29 and 0.35) (Table 4). The confidence interval for the associations between TMT and general cognitive function, and between TMT and processing speed did not overlap with the confidence interval for the association between TMT and memory.

**Table 3.** Genetic correlations (standard errors) using GCTA-GREML (under diagonal) and LD score regression (above diagonal) for the trail making tests and verbal-numerical reasoning in UK Biobank.

|  | TMT A | TMT B | TMT B-A | VNR |
| --- | --- | --- | --- | --- |
| TMT A | - | 0.921 (0.07) | 0.827 (0.17) | 0.558 (0.11) |
| TMT B | 0.840 (0.09) | - | 0.981 (0.03) | 0.767 (0.07) |
| TMT B-A | 0.640 (0.19) | 0.958 (0.03) | - | 0.793 (0.10) |
| VNR | 0.589 (0.36) | 0.636 (0.14) | 0.587 (0.18) | - |

TMT A, trail making part A; TMT B, trail making part B; TMT B-A, trail making part B minus trail making part A; VNR, verbal-numerical reasoning.

**Table 4.** Genetic correlations (standard error), derived using LD score regression, between three trail making test in UK Biobank and general cognitive function, processing speed, and memory from the CHARGE consortium. TMT A, trail making part A; TMT B, trail making part B; TMT B-A, trail making part B – part A. Tests that survived FDR correction (p = 0.0404) are shown in bold.

|  | TMT A |  |  | TMT B |  |  | TMB B-A |  |  |
| --- | --- | --- | --- | --- | --- | --- | --- | --- | --- |
| | $r_g$ | SE | p | $r_g$ | SE | p | $r_g$ | SE | p |
| General cognitive function | 0.6076 | 0.11 | **$2.73 \times 10^{-8}$** | 0.6951 | 0.07 | **$3.02 \times 10^{-22}$** | 0.6566 | 0.08 | **$9.84 \times 10^{-15}$** |
| Processing speed | 0.7614 | 0.15 | **$3.01 \times 10^{-7}$** | 0.7646 | 0.08 | **$7.69 \times 10^{-20}$** | 0.6877 | 0.10 | **$8.36 \times 10^{-13}$** |
| Memory | 0.2945 | 0.14 | **0.0404** | 0.3532 | 0.09 | **0.0001** | 0.2994 | 0.11 | **0.0055** |

### Polygenic prediction

Polygenic profiles based on the TMT A summary results from the CHARGE consortium significantly predicted all three trail making tests in UK Biobank (*β* between 0.016 and 0.029, Table 5). Polygenic profiles based on the TMT B summary results from the CHARGE consortium significantly predicted all three trail making tests in UK Biobank (*β* between 0.024 and 0.036, Table 5). The strongest associations for the other cognitive test polygenic profiles (general cognitive function, processing speed, and memory) were found for general cognitive function polygenic profiles predicting TMT B in UK Biobank, explaining 0.67% of the variance. The full results including all thresholds can be found in Supplementary Table 2a.

**Table 5.** Associations between polygenic profiles based on CHARGE cognitive tests and trail making in UK Biobank. FDR correct p-value = 0.04455473. Associations are scored such that a higher polygenic score indicates better performance on the TMT tests.

|  |  | Phenotypes in UK Biobank |  |  |  |  |  |  |  |  |  |  |  |
| --- | --- | --- | --- | --- | --- | --- | --- | --- | --- | --- | --- | --- | --- |
|  |  | Trail making test part A |  |  |  | Trail making test Part B |  |  |  | Trail making test part B – part A |  |  |  |
|  |  | best threshold | beta | r2 % | p | best threshold | beta | r2 | p | best threshold | beta | r2 | p |
| Polygenic profiles | Trail making test part A | 0.5 | 0.0211 | 0.04 | **0.0006** | 0.5 | 0.0288 | 0.08 | **1.34×10<sup>-6</sup>** | 0.5 | 0.0160 | 0.03 | **0.0108** |
|  | Trail making test Part B | 1 | 0.0274 | 0.08 | **7.36×10<sup>-6</sup>** | 0.5 | 0.0355 | 0.13 | **2.07×10<sup>-9</sup>** | 0.1 | 0.0239 | 0.06 | **0.0001** |
|  | General cognitive function | 0.5 | 0.0583 | 0.30 | **2.44×10<sup>-19</sup>** | 0.5 | 0.0867 | 0.67 | **3.11×10<sup>-43</sup>** | 0.5 | 0.0649 | 0.38 | **1.49×10<sup>-22</sup>** |
|  | Processing speed | 0.5 | 0.0478 | 0.22 | **1.45×10<sup>-14</sup>** | 0.5 | 0.0771 | 0.57 | **2.60×10<sup>-37</sup>** | 0.5 | 0.0616 | 0.37 | **4.34×10<sup>-22</sup>** |
|  | Memory | 0.05 | 0.0200 | 0.04 | **0.0011** | 1 | 0.0343 | 0.12 | **9.31×10<sup>-9</sup>** | 1 | 0.0298 | 0.09 | **2.32×10<sup>-6</sup>** |

The GWAS results for the three trail making phenotypes in UK Biobank were used to create polygenic profile scores in two independent cohorts; Generation Scotland (GS), and the Lothian Birth Cohort of 1936 (LBC1936) (Figure 3 and Supplementary Tables 2b and 2c). The polygenic profiles for TMT A significantly predicted digit-symbol substitution, verbal fluency, Mill Hill vocabulary, fluid cognitive function, and general cognitive function in GS (β between 0.018 and 0.039). In the LBC1936, the polygenic profiles for TMT A significantly predicted trail making part B (β = 0.102), fluid cognitive function, memory and change in fluid cognitive function (β between 0.079 and 0.094). Significant predictions were observed across almost all thresholds for TMT B and TMT B-A for the cognitive phenotypes measured in GS. In LBC1936, polygenic profiles for TMT B and TMT B-A were both significantly associated with fluid cognitive function, processing speed, cognitive function at age 11 and cognitive function at age 70. TMT B was also significantly associated with trail making part B, memory and change in fluid cognitive function between age 11 and age 70. The strongest association in GS was found between the polygenic profile for TMT B and Wechsler digit symbol substitution; this association explained 0.53 % of the variance, at a SNP inclusion threshold of all SNPs from the GWAS. The largest proportion of variance explained in LBC1936 was 1.75 % for trail making part B using the TMT B polygenic score with a SNP inclusion threshold of all SNPs from the GWAS. The complete results can be found in Supplementary Tables 2b and 2c.

**Figure 3.**
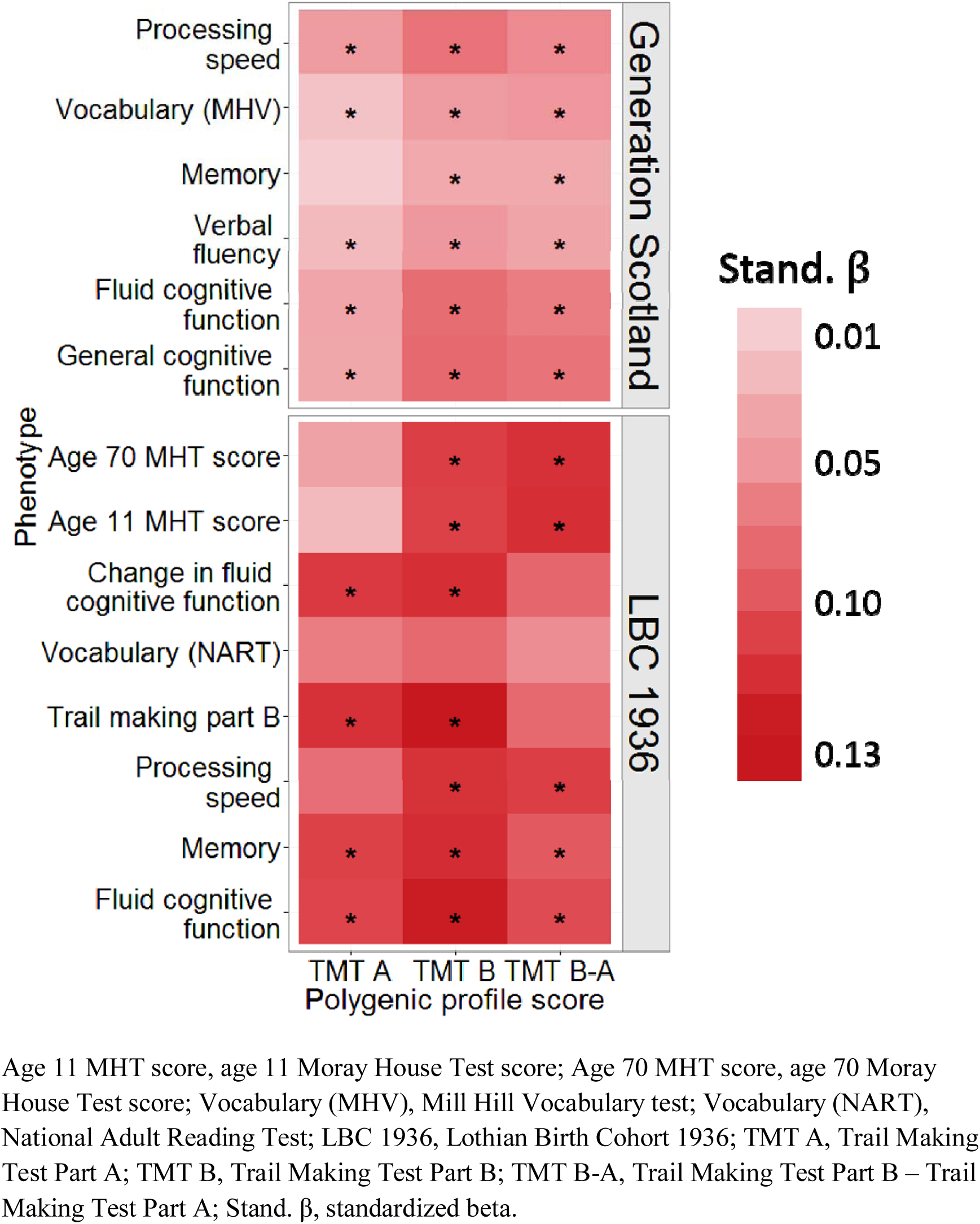
Heat map of associations between the polygenic profile scores for trail making in UK Biobank and cognitive function in Generation Scotland and the Lothian Birth Cohort 1936 (LBC 1936). * indicates FDR corrected significant associations (FDR corrected p-value <= 0.029 (Generation Scotland) or 0.025 (LBC 1936)). Further information can be found in Supplementary Tables 2b and 2c.

### Meta-analysis

In the meta-analysis of the combined dataset (UK Biobank and CHARGE, combined N TMT A = 29 251, combined N TMT B = 30 022), no genome-wide significant SNPs were observed for TMT A or TMT B (Supplementary Figure 3).

## Discussion

This study finds new genome-wide significant variants associated with trail making test performance, and provides the first SNP-based heritability estimates for the three widely-used TMT measures. We identified high genetic correlations between trail making, verbal-numerical reasoning, general cognitive function, and processing speed, and somewhat lower genetic correlations with memory. Using only common SNPs to create a polygenic score for TMT B, we were able, at best, to predict ~2 % of the variance in trail making part B in LBC1936. Taken together, these analyses point to a considerable degree of shared polygenic architecture for TMT performance with general cognitive function and processing speed measures, in particular. The results are consistent with the hypothesis that trail making is genetically and phenotypically similar to general fluid cognitive function ^9, 20, 34^.

Univariate GCTA-GREML analyses suggested SNP-based heritability estimates for TMT A, TMT B and TMT B-A of 8%, 22% and 18% respectively. These provide a first estimate of the contribution of common SNPs to the phenotype of trail making and suggest that common SNPs account for around half of the additive genetic variation for trail making, based on twin and family studies ^30^. SNP-based heritability studies for other complex traits also report similar differences between SNP based and pedigree based heritability ^48, 66^. Twin and family studies estimate heritability based on all causal variants, both common and rare, whereas SNP-based studies estimate heritability only on genotyped SNPs in LD with causal variants. It is possible that causal variants have lower minor allele frequencies than the genotyped SNPs, leading to incomplete LD between unknown causal variants and genotyped SNPs ^67^.

Previous studies of the genetic overlap between trail making and other cognitive abilities have shown genetic overlap between trail making and general cognitive function using a twin design ^30, 34^. Our results add to this by using a molecular genetic design and showing shared genetic aetiology between trail making and verbal numerical reasoning in UK Biobank, as well as with general cognitive function, processing speed, and memory from the CHARGE consortium. The estimates of the genetic correlations within UK Biobank were similar for both GCTA-GREML and LDS regression, suggesting that these results are unlikely to constitute false positives.

We also used polygenic profile scores to estimate the genetic overlap between TMT and other cognitive abilities. First we created polygenic profiles based on the CHARGE consortium summary GWAS data (including trail making, general cognitive function, processing speed, and memory) and found that the polygenic profile for general cognitive function was the best predictor of all TMT scores in UK Biobank compared to either of the other polygenic profiles. All estimates were small (< 1% of variance was accounted for), but should be considered the minimum estimate of the variance explained. The polygenic profile method prunes SNPs in LD and assumes that a single causal variant is tagged in each LD block. If that assumption is violated, the proportion of variance explained will be underestimated. Nevertheless, the unequal sample sizes from which the polygenic profile scores were derived (TMT ~ 6000; general cognitive function ~54 000; processing speed ~ 32 000, memory ~ 29 000) may have led to an underestimation of the ability of CHARGE TMT score to predict UK Biobank TMT test performance.

Our combined meta-analysis of UK Biobank and CHARGE trail making did not yield any significant SNPs. This could be ascribed to the following factors. The SNPs for the CHARGE consortium are imputed to the HAPMAP 2 reference panel (N ~ 2M SNPs) whereas the UK Biobank sample is imputed to a combination of the UK10 haplotype and the 1000G reference panel (N ~ 17M SNPs). For TMT A, 2 332 746 SNPs overlapped between UK Biobank and CHARGE, and for TMT B, 2 466 810 SNPs overlapped between the two samples. The SNPs that reached genome-wide significance in the UK Biobank GWAS had low minor allele frequencies and therefore likely did not pass quality control (MAF > 0.01) in the smaller CHARGE consortium sample (N~6000), this smaller sample size limits statistical power to detect genome-wide significant variants.

In addition to those discussed above, our study has other limitations. The measure of fluid cognitive function provided by UK Biobank (which we have called verbal-numerical reasoning) showed a relatively modest age-related trajectory, in contrast to the steeper and well-replicated age-related decline that would be expected for this construct ^68, 69^ (Supplementary Figure 4). This may partly explain the relatively modest correlation of verbal-numerical reasoning with TMT performance, when compared to those previously reported ^9^ and may have also had a bearing on our analyses of genetic overlap within UK Biobank. In addition, the UK Biobank sample did not have sufficient breadth of contemporaneously-administered standardised/validated cognitive tests to be able to construct a robust measure of general cognitive function (such as in other large samples^9, 20, 70^). This limited our ability to perform a more detailed analysis of the phenotypic or genetic overlap between trail making and general cognitive function in UK Biobank itself.

This study has several strengths. It has the largest single sample size to date of a GWAS for trail making, offering greater statistical power while excluding bias caused by sample or phenotypic heterogeneity. Population stratification was minimized by only using individuals of white British ancestry. This study shows the first estimates of the heritability for TMT using molecular genetic data, uses a comprehensive battery of techniques to examine genetic architecture of TMT, and used the same trail making test and administration protocol across the whole sample. The use of both GWAS summary data from the CHARGE consortium, and the prediction of the UK Biobank TMT GWAS into GS and LBC1936 cognitive phenotypes allowed a detailed examination of the shared genetic aetiology between TMT and cognitive phenotypes.

These strengths have enabled a detailed characterisation of the shared genetic aetiology between performance on TMT and other cognitive abilities. Our results, spanning methodologies and cohorts, provide strong evidence for a shared genetic aetiology between TMT performance and general cognitive function and processing speed, which are themselves strongly phenotypically and genetically correlated. When the full genetic data from UK Biobank on half a million individuals becomes available, it would enable robust replication and extensions of the current findings. More detailed cognitive testing is planned for UK Biobank, and these data can be used in future studies to further examine the genetic overlap between TMT and other cognitive functions.

## Acknowledgments

This research was conducted using the UK Biobank Resource. The work was undertaken in The University of Edinburgh Centre for Cognitive Ageing and Cognitive Epidemiology, part of the cross council Lifelong Health and Wellbeing Initiative (MR/K026992/1); funding from the BBSRC and Medical Research Council (MRC) is gratefully acknowledged. The Lothian Birth Cohort is supported by Age UK (Disconnected Mind Project). AMM and IJD are supported by a Wellcome Trust Strategic Award (104036/Z/14/Z).

### Conflict of Interest

IJD is a participant in UK Biobank.

